# Activation of mGlu2 receptors reverses persistent post-methamphetamine deficit in object-in-place recognition memory

**DOI:** 10.64898/2026.05.25.727633

**Authors:** Viktoria Galbava, Lizhen Wu, Marek Schwendt

## Abstract

**Background/Objectives:** Persistent cognitive impairments are prevalent in methamphetamine (meth) use disorder and contribute to maladaptive decision-making and increased relapse vulnerability. There are currently no effective treatments for meth-associative cognitive deficits, and their neurobiological underpinnings remain incompletely understood. This study investigated the effects of chronic meth self-administration on episodic-like recognition memory and evaluated whether pharmacological potentiation of metabotropic glutamate receptor subtype 2 (mGlu2) could rescue these deficits.

**Methods:** Adult male Sprague-Dawley rats underwent 7 days of limited- (1h/day) followed by 14 days of extended-access (6h/day) meth self-administration, followed by 30 days of abstinence. Recognition memory was assessed using the object-in-place (OIP) task. A positive allosteric modulator of mGlu2 receptors, LY-487379 (25 mg/kg, s.c.), was administered prior to the memory test. In parallel, changes in total and surface mGlu2/3 protein levels in the prelimbic and perirhinal cortices were evaluated.

**Results:** Rats with extended access to meth self-administration exhibited escalated drug intake and persistent deficits in OIP memory. Administration of LY-487379 reversed this deficit. Total mGlu2/3 protein levels were unaltered; however, meth exposure was associated with a significant increase in surface mGlu2/3 receptor expression in both cortical regions examined.

**Conclusions:** These results demonstrate that chronic meth produces persistent cognitive dysfunction that can be rescued by mGlu2 receptor potentiation. The observed increase in surface mGlu2/3 expression may represent a compensatory response to chronic glutamatergic dysregulation, but it appears to be insufficient to restore cognitive function alone, without pharmacological enhancement. The current data encourage further exploration of mGlu2 role in stimulant-associated cognitive dysfunction.

**Highlights:**

Chronic methamphetamine self-administration produced persistent deficits in episodic-like recognition memory in male rats and dysregulation of mGlu2/3 receptors in the prelimbic and perirhinal cortices.
Systemic pharmacological potentiation of mGlu2 receptors rescued meth-associated memory deficits.
mGlu2 receptor potentiation may represent a promising therapeutic strategy for treating stimulant-associated cognitive dysfunction.
Increased surface mGlu2/3 expression may represent a compensatory adaptation to post-methamphetamine glutamatergic dysfunction, but it is not sufficient to restore cognition alone, without pharmacological enhancement.

## 1. Introduction

Methamphetamine (meth) is a highly addictive synthetic amphetamine-type stimulant. As a result, repeated exposure to meth can rapidly progress from controlled recreational to uncontrolled, compulsive use and ultimately lead to meth use disorder (MUD).

Problematic meth use and MUD remain a major public health challenge in the United States and worldwide, driven by a rising number of users (recently estimated at 2.4 million in the US and 316 million globally) [1,2], and a high prevalence of MUD among the user population [3]. These challenges are compounded by the absence of effective pharmacotherapies for MUD, and by the high burden of meth-associated complications, such as overdose, as well as cardiovascular, neurological, and psychiatric complications [4–6]. Despite these adverse consequences, heavy meth users often struggle to maintain abstinence due to heightened drug craving and cognitive deficits that persist long after the cessation of meth use [7]. Additionally, due to a lack of pharmacotherapies for MUD, the current treatment primarily relies on behavioral interventions, such as cognitive behavioral therapy (CBT) [8], of which the moderate efficacy is further compromised by the presence of meth-induced cognitive deficits. Emerging evidence suggests that the manifestation of craving and cognitive deficits in MUD follows a similar trajectory, and these two phenomena are also likely cross-dependent [9,10]. Therefore, it has been suggested that understanding and reversing cognitive deficits in MUD would also improve primary treatment outcome (12-month abstinence rate) [11].

The majority of clinical research finds that prolonged and heavy meth use is associated with mild-to-severe impairments across multiple cognitive domains, including learning and memory, executive functioning, attention, and information processing speed [7,12–16]. Additionally, episodic memory problems are frequently observed in subjects with MUD [7,12,17], and can significantly contribute to increased relapse risk [18]. As studies in human subjects cannot readily establish whether episodic memory deficits stem from the direct disruption of brain circuits supporting episodic memory or, indirectly, from executive or attentional dysfunction [19], preclinical research is needed to provide mechanistic and functional insights into episodic memory deficits. Our previous work has demonstrated that extended access to self-administered meth and a non-contingent meth binge regimen in rats produce deficits in recognition memory, as assessed using two validated tasks, novel object recognition (NOR) and object-in-place (OIP) recognition [20–22]. These tasks are thought to probe episodic-like memory by requiring animals to recall both the identity (“what”) of objects and their spatial context (“where”). Of the two, the OIP task represents a more demanding version of associative recognition memory that depends on synchronous activation of circuits encompassing the perirhinal (PrH) and prefrontal (prelimbic, PrL) cortices (as well as the hippocampus) [23,24]. More recent work showed that the PrH-to-PrL circuit is necessary for the expression of recognition memory, as chemogenetic inhibition abolishes recognition memory, while chemogenetic activation of PrH-to-PrL projection rescues meth-induced recognition memory deficits [25]. In addition to recognition memory deficits, prior work from our group (and our collaborators) found that a history of extended access to meth self-administration is associated with dysregulation of glutamatergic synaptic plasticity and altered expression of select ionotropic and metabotropic glutamate receptors in the PrL and PrH [14,21,26,27]. However, this body of work focused on the period of early abstinence, 7-14 days after discontinuation of meth self-administration. Consequently, it remains unclear whether episodic-like memory deficits persist or gradually recover with prolonged abstinence. Clinical evidence suggests that while some cognitive deficits recover over time, others can persist for months or longer [28,29].

Here, we addressed this question by investigating: (a) whether OIP memory deficits persist for up to 30 days after discontinuation of extended access meth self-administration, and (b) whether systemic allosteric activation of metabotropic glutamate subtype 2 (mGlu2) receptors with LY-487379 rescues such OIP memory deficits. The choice of mGlu2 as a target of choice is based on prior work, suggesting the allosteric activation of mGlu2 receptors reduces persistent meth seeking [30], enhances cognitive flexibility in naïve rats [31], and reverses recognition memory deficits in experimental models of psychosis [32,33]. Finally, we propose to explore whether post-meth deficits in episodic-like memory are accompanied by changes in the active pool of mGlu2 receptors by quantifying cell-surface mGlu2 receptors in both brain regions of interest.

Prior work suggests that mGlu2 (or mGlu/23) receptors in rodent frontal cortex undergo changes with chronic meth exposure [34–37].

## 2. Materials and Methods

### Subjects

Adult male Long-Evans rats (N=43, Charles-River Laboratories, 275-300g upon arrival) were used in this study. The sample size and animal numbers were determined by power analysis of our prior data [21,22,34]. The rats were individually housed in a climate-controlled vivarium on a 12-hour reversed light-dark cycle (dark phase: 7 AM – 7 PM). The single housing of animals was necessitated by the implantation of an external catheter and harness for a standard i.v. drug self-administration (IVSA) procedure. All behavioral testing was conducted during the dark phase of the cycle. Rats received ad libitum access to standard rat chow and water upon arrival and during acclimation. At the start of behavioral testing and throughout the remainder of the experiment, rats were maintained under mild food restriction (85% of their ad libitum intake). All animal procedures were conducted and approved by the Medical University of South Carolina Institutional Animal Care and Use Committee (IACUC). All procedures complied with the Guide for the Care and Use of Laboratory Animals (National Research Council (US) Committee for the Update of the Guide for the Care and Use of Laboratory Animals 2011).

### Pharmacological agents

(+)-Methamphetamine hydrochloride (meth) was obtained through the National Institute of Drug Abuse (NIDA) drug supply program. Meth was dissolved in 0.9% sterile saline to attain a concentration of 0.40 mg/ml. All rats received a dose of 0.1 mg/kg/infusion during IVSA. Selective positive allosteric modulator of mGlu2 receptors: N-(4-(2-methoxyphenoxy)phenyl)-N-(2,2,2-trifluoroethylsulfonyl)pyrid-3-ylmethylamine (LY-487379 hydrochloride) was purchased from Tocris/Bio-techne (Minneapolis, MN, USA; Purity: ≥98%; cat. No.3283). LY-487379 is a highly specific, brain-penetrant mGlu2 receptor positive allosteric modulator (PAM) with documented pro-cognitive properties [38–40]. LY-487379 was dissolved in 0.5 % carboxymethylcellulose / 0.5 % Tween-80 / sterile saline, incubated at 50 °C for 15 minutes, and sonicated to aid LY-487379 solubilization. The mixture of 0.5 % carboxymethylcellulose / 0.5 % Tween-80 / sterile saline served as a vehicle. The dose of LY-487379 used in this study was carefully selected based on previous studies attesting to behavioral efficacy and lack of overt side effects [40–43].

### Surgery

Rats received chronically indwelling catheters into the right jugular vein as previously described [43]. Briefly, rats were implanted with intravenous catheters under ketamine (87.5 mg/kg i.p.) and xylazine (5 mg/kg i.p.) anesthesia, with ketorolac (2.0 mg/kg i.p.) administered for analgesia. A Silastic catheter was inserted into the right jugular vein, secured with silk sutures, and routed subcutaneously to exit between the scapulae, where it was connected to a stainless-steel cannula within an infusion harness to allow repeated intravenous drug delivery. Postoperatively, rats received cefazolin (10 mg/0.1 ml) to prevent infection, and catheters were maintained with heparinized saline (10 U/ml). Before each self-administration session, catheters were flushed with heparinized saline; after each session, they were flushed with cefazolin followed by heparinized saline to maintain patency and sterility. Catheter patency was periodically verified using intravenous methohexital sodium (10 mg/ml in 0.9% NaCl), which induces rapid loss of muscle tone when the catheter is functional.

### Meth self-administration and abstinence

Following at least five days of post-surgical recovery, rats were randomly assigned to either the meth self-administration or the control groups. Meth self-administration occurred in standard operant chambers (Med Associates, St Albans, VT) equipped with active and inactive levers, cue lights, and a tone generator. Rats received intravenous meth (20 μg/50 μl over 2 s) under an FR1 schedule, where each active lever press triggered an infusion paired with a 5-s tone (78 dB, 4.5 kHz) and light cue, followed by a 20-s timeout. Training consisted of 7 daily limited-access (1h) sessions and 14 daily extended-access (6h) sessions. Inactive lever presses and responses during timeout were recorded but had no consequences. Concurrently, control animals received catheter surgery and underwent a yoked saline procedure (with infusions and cue-presentations matching the meth ‘partner’ rat) or were assigned to a ‘playback’ condition and received matching presentations of drug-associated cues. Catheter patency was regularly checked using methohexital sodium (10 mg/ml in 0.9% saline; Eli Lilly, Indianapolis, IN, USA). All operant sessions occurred during the dark phase. Three rats were excluded from analysis for jugular catheter failure or illness during IVSA training (n = 2 from meth IVSA group; n = 1 from control group). Next, rats underwent 30 days of home-cage abstinence with daily handling. See Figure 1 for the timeline of the experiment.

**Figure 1.**
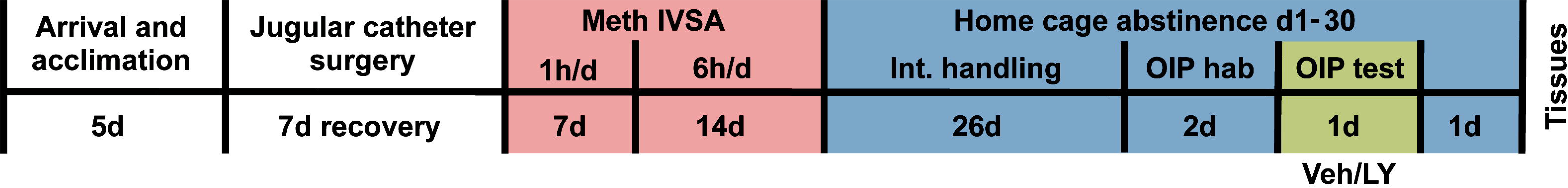
Experimental Timeline. Upon arrival, rats were habituated in their home cages for 5 days, followed by surgery and a 7-day recovery period. Meth intravenous self-administration (IVSA) consisted of short-access sessions (1 h/day) for 7 days, followed by long-access sessions (6 h/day) for 14 days using an FR1 schedule of reinforcement. Next, rats underwent home-cage abstinence, during which the OIP task performance was tested on days 27-29.

### Object-in-Place memory task

Recognition memory testing using the object-in-place task was conducted as described previously [45,46]. A subgroup of rats (N = 32) was used for the OIP testing. Briefly, rats were tested on days 28-30 after the end of meth self-administration, using an elevated round platform (125 cm in diameter) as an OIP test arena. On days 28-29, rats received a daily habituation session during which they were placed in the arena with no objects and left undisturbed to explore for 5 minutes. On day 30, during the familiarization phase (or sample phase), rats were placed in the apparatus for 5 min with 4 distinct objects positioned in adjacent corners. Sixty minutes after object familiarization (and 30 minutes prior to the OIP test), rats received a single subcutaneous injection of LY-487379 25 mg/kg, or vehicle (see the Pharmacological agents section for the vehicle description).

Animals were randomly assigned to the Veh vs LY group, counterbalanced for between-group meth intake. The selection of objects, the timing of the injection, and the dose were carefully chosen based on previous studies [41,42,47]. An OIP memory test was conducted by placing the rat in the apparatus for 3 min with the same objects, except that the position of two objects was changed (“same” and “changed”, respectively).

Object placement was counterbalanced. Behavior was recorded and stored using Noldus tracking software (EthoVision XT 6.0), and an observer naive to the experimental conditions scored behavior. Object exploration was defined as sniffing or touching the object with the nose, but not sitting, leaning, or standing on objects. Animals were euthanized and tissues collected 24 hours after the OIP test. See Figure 1 for the detailed timeline of the experiment.

### Slice preparation of surface biotinylation

Surface biotinylation of acute brain slices was performed as described previously, with some modifications [48–50]. After rapid decapitation, rat brains were removed and briefly chilled in ice-cold ACSF oxygenated with 95% O2/5% CO2. Next, 2 mm-thick coronal slices containing the PFC (AP +3.00 to +4.20 mm from bregma) and PrH AP (−3.24 to −5.16 mm from bregma) were prepared using the rat brain matrix (ASI instruments, Warren, MI, USA). The tissues were then bilaterally dissected using a 2-mm micropunch (Harris-Unicore, Ted Pella, Redding, CA, USA) and cut into 250 μm-thick mini-sections using a McIlwain Tissue Chopper (Stoelting, Wood Dale, IL, USA). Next, acute sections were biotinylated with 1 mg/ml EZ link NHS-SS-Biotin/ACSF (Thermo Fisher Scientific, Rockford, IL, USA) for 1 hr at 4°C. The biotinylation reaction was terminated with a 100 mM glycine/ACSF buffer. Biotinylated slices were lysed by brief sonication in lysis buffer (25 mM HEPES, 150 mM NaCl, 1% Triton X-100, 0.1% SDS) supplemented with protease and phosphatase inhibitors and incubated on ice for 15 mins. Insoluble debris was removed by centrifugation. An equal aliquot of pre-cleared solubilized total protein fraction (T) was set aside and frozen at −80°C. Biotinylated proteins were captured by incubating with Neutravidin agarose beads (Thermo Fisher Scientific) overnight at 4°C.

After non-biotinylated, intracellular (I) proteins were removed, bound biotinylated surface (S) proteins were recovered from the beads using an elution buffer (62.5 mM Tris-HCl, pH 6.8, 20% glycerol, 2% SDS, and 50 mM dithiothreitol, pH 6.8) and incubated at 95°C for 10 min. Protein concentrations were determined using the BCA assay (Thermo Fisher Scientific), and proteins of interest were analyzed by immunoblotting as described below. See Figure 4 for more details.

### Immunoblotting

Brain-region-specific levels of candidate proteins were assessed by immunoblotting in individual fractions (total lysate, intracellular, and surface) obtained as described in Section 2.6 and previously [48]. A subgroup of 24 rats from Ctr-Veh (n = 12), Meth-Veh (n = 7), and Meth-LY (n = 5) was used for the tissue analysis. Briefly, equal amounts of protein from each fraction were separated on 4–15% SDS–polyacrylamide gels and transferred to PVDF membranes. To reduce non-specific binding, membranes were incubated in 5% milk in Tris-buffered saline containing 0.1% Tween-20 (blocking buffer) for 1 h at room temperature. Next, membranes were incubated overnight at 4 °C with primary antibodies diluted in blocking buffer. The following primary antibodies and dilutions were used: mGlu2/3 rabbit polyclonal (Upstate, #06-676, 1:15,000; anti-rabbit secondary 1:15,000), RanBP9 rabbit polyclonal (Abcam #ab140627, 1:10,000; anti-rabbit 1:10,000), Gαi mouse monoclonal (New East Biosciences, #26003, 1:10,000; anti-mouse 1:10,000), β-tubulin rabbit polyclonal (Abcam #AB6046, 1:80,000; anti-rabbit 1:80,000), and syntaxin-1a rabbit polyclonal (Abcam #ab41453, 1:40,000; anti-rabbit 1:40,000). Note that separate analysis of mGlu2 and mGlu3 receptor expression was not pursued in this study, due to the inconsistent signal obtained with the mGlu2-specific antibody (Millipore-Sigma #07-261-I). Membranes were incubated with appropriate HRP-conjugated secondary antibodies, and bands were detected using enhanced chemiluminescence (ECL Plus) and captured on a sensitive X-ray film (Hyperfilm, GE Healthcare). Membranes were stripped and re-probed with housekeeping markers (β-tubulin) or membrane-specific markers (syntaxin-1a) to control for loading, transfer efficiency, and surface biotinylation specificity. Band intensity was quantified as integrated density using ImageLab software (Bio-Rad, Hercules, USA). Immunoblotting data analysis was performed with experimenter blinding.

### Statistical analysis

Meth intake (mg/kg) and lever responding served as the primary dependent variables and were evaluated across 14 and 21 days of self-administration sessions, respectively, using a one-way repeated-measures ANOVA, followed by Geisser-Greenhouse correction when appropriate. For the OIP task, the main outcome measure was the amount of time animals spent exploring each object. Exploration time for the two objects that remained in their original positions was combined, and the same procedure was applied to the two objects that were relocated. These values were then used to calculate a recognition index defined as exploration of the moved objects divided by the total exploration of moved plus stationary objects. Recognition index values were analyzed using a two-way ANOVA. Between-group comparisons (meth vs. control) for behavioral and biochemical endpoints were performed with independent (two-sample) t-tests.

Integrated density values for target proteins were first normalized to those of control markers (syntaxin-1a or β-tubulin), expressed as per cent values relative to controls (percent of control), and analyzed with the same independent-samples t-test approach. Outliers were identified and excluded using the ROUT method, with a false discovery rate of Q = 1%. All results are reported as mean ± SEM, and statistical significance was assessed at α = 0.05. Random assignment of animals and experimenter blinding were implemented for immunoblotting, functional testing, and outcome assessment throughout the study.

## 3. Results

### Meth IVSA

Rats consistently discriminated between the active and inactive levers across all 21 days of IVSA, and as expected, selectively increased their active lever responding with the onset of the extended (6-hour) access. Two-way repeated-measures Anova revealed a significant main effect of Day (F 5.548, 164.8 = 19.92, p < 0.0001), a significant main effect of Lever (F 1, 30 = 79.80, p < 0.0001), and a significant Day × Lever interaction (F 5.548, 164.8 = 15.02, p < 0.0001). Tukey’s post hoc comparisons confirmed a significant difference between active and inactive lever responding on each day of IVSA (Figure 2a). Analysis of the daily amount of self-administered meth (in mg/kg) revealed a significant escalation of drug intake over the 6-hour access period (F 4.765, 70.75 = 4.794, p < 0.001). Dunnett’s multiple comparisons indicated that intake on days 15 and 18-21 was significantly higher than the average starting extended-access intake (days 8–10) (Figure 2b). Simple start-to-end analysis of extended access meth intake confirmed a significant escalation between the initial (days 8–10) and the final (days 19–21) intake [two-tailed paired t-test, t(15) = 5.288, p < 0.0001] (Figure 2c). Overall mean meth intake in all rats was 61.52 ± 5.13 mg/kg. There were no differences in meth intake between rats later assigned to LY-487379 or the vehicle group [two-tailed paired t-test, t(14) = 1.019, p = > 0.05, ± SEM = 11.26 ± 11.05].

**Figure 2.**
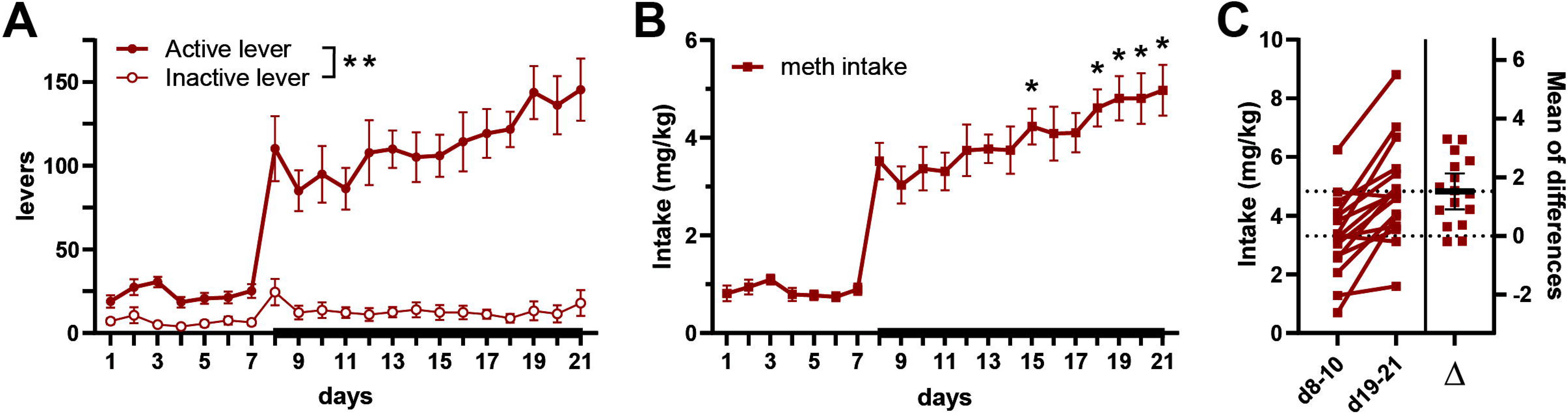
Intravenous Meth self-administration. (a) Active lever responding intensified over time, with a marked increase after day 8 (beginning of the 6h/day extended meth access), whereas inactive lever responding remained low. There were significant effects of Day [F 5.548, 164.8 = 19.92, p < 0.0001], Lever [F 1, 30 = 79.80, p < 0.0001], and a Day × Lever interaction [F 5.548, 164.8 = 15.02, p < 0.0001]; (b) Meth intake (mg/kg) across 21 daily sessions. Intake was low during days 1–7 of limited 1h/day access and increased with extended 6h/day access, followed by progressive intake escalation. There was a significant effect of Day [F 4.765, 70.75 = 4.794, p < 0.001], with post-hoc analysis revealing that intake significantly escalated on days 15, 18–21 (*p < 0.05–0.001; Dunnett’s test) vs. the initial day 8-10.; (c) Estimation plot showing extended access intake (mg/kg) during days 8–10 vs. days 19–21. Each pair of points connected by lines represents an individual subject (n = 16). Intake increased from d8–10 to d19–21. The right panel displays the paired mean difference (Δ) with individual values shown. Statistical analysis using a two-tailed paired t-test revealed a significant increase in intake over time [t(15) = 5.288, p < 0.0001]. Groups: Meth (n = 16). Data represent the mean ± SEM.

### OIP memory task

Rats were tested on the OIP task on day 28 of abstinence following two days of habituation to the apparatus. Initial exploration of the four objects did not differ during the familiarization/sample session (mean seconds ± SEM; plastic bottle, 21.71 ± 2.28; light bulb, 18.40 ± 1.40; paint roller, 19.5 ± 2.02; PVC pipe, 22.01 ± 1.89). Testing occurred 30 min after administration of LY-487379 (25 mg/kg) or vehicle, and 90 min after the sampling phase, as shown in Figure 3a. For the OIP test, a two-way ANOVA revealed a significant main effect of Treatment [F 1,28 = 4.342, p < 0.05], indicating that LY-487379 treatment significantly influenced recognition performance across conditions. In addition, there was a significant Drug × Treatment interaction [F 1,28 = 4.964, p < 0.05], suggesting that the effect of LY-487379 depended on prior meth exposure. Post hoc Tukey’s multiple comparisons showed that rats with a history of meth (and treated with vehicle) exhibited impaired recognition memory compared to vehicle-treated controls (p < 0.05), suggesting persistent meth-induced memory impairment. And further, meth rats treated with LY-487379 showed significantly improved recognition memory compared to meth-vehicle counterparts (p < 0.05). In fact, recognition memory performance of meth+LY rats was comparable to that of the control+LY group (p = 0.9762). LY-487379 administration produced no further enhancement of recognition memory in control animals (p = 0.9996), consistent with a treatment ‘ceiling effect’ in unimpaired animals.

**Figure 3.**
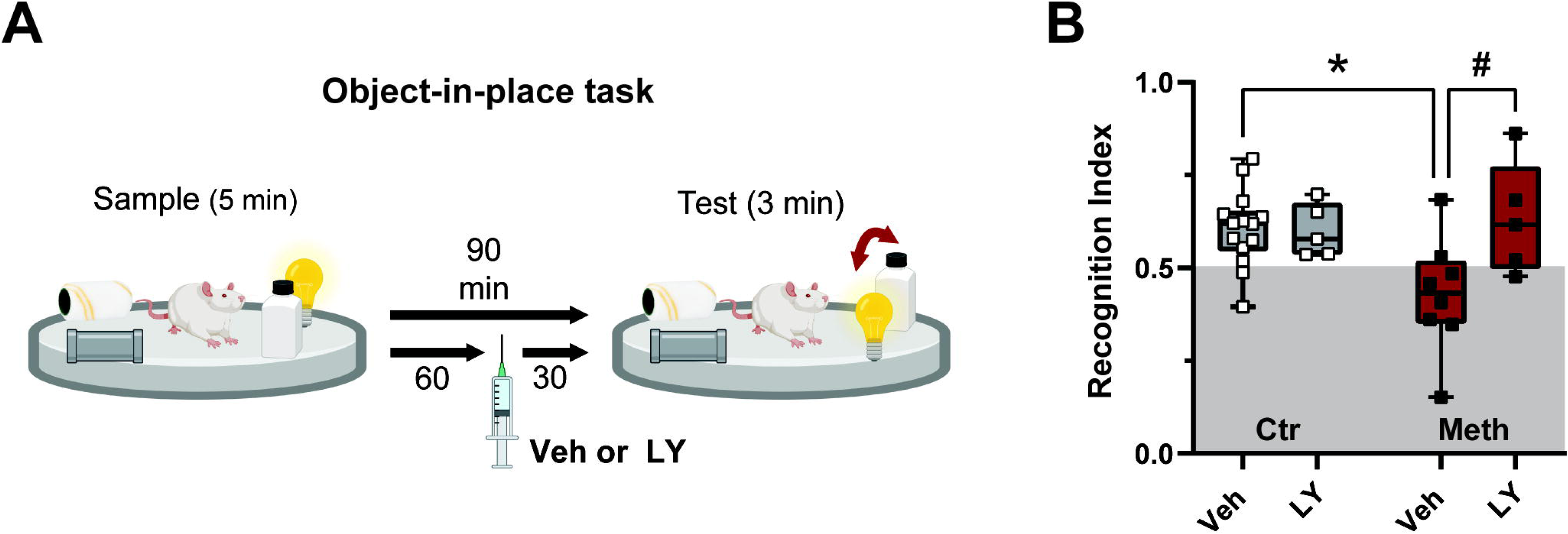
Object-in-place recognition memory task and the effects of mGlu2 PAM treatment. (a) During the sampling phase, rats were placed in the arena for 5 min with four distinct objects positioned in adjacent corners. Sixty minutes after familiarization (and 30 min prior to testing), rats received a single injection of LY-487379 (25 mg/kg s.c.) or vehicle. OIP memory was then assessed during a 3 min test session for which two of the objects were relocated while the remaining objects stayed in their original positions; (b) Recognition index in control (Ctr) and meth self-administering (Meth) rats treated with vehicle (Veh) or LY-487379 (LY). Two-way ANOVA revealed a significant main effect of Treatment [F 1,28 = 4.342, p < 0.05] and interaction [F 1,28 = 4.964, p < 0.05]. Follow-up pairwise comparisons revealed that meth exposure reduced recognition performance compared to controls (*p < 0.05), an effect reversed by LY treatment (#p < 0.05). LY administration had no effect in control rats. Groups: Ctr-Veh (n = 14), Ctr-LY (n = 5), Meth-Veh (n = 8), Meth-LY (n = 5). Data represent the mean ± SEM.

**Figure 4.**
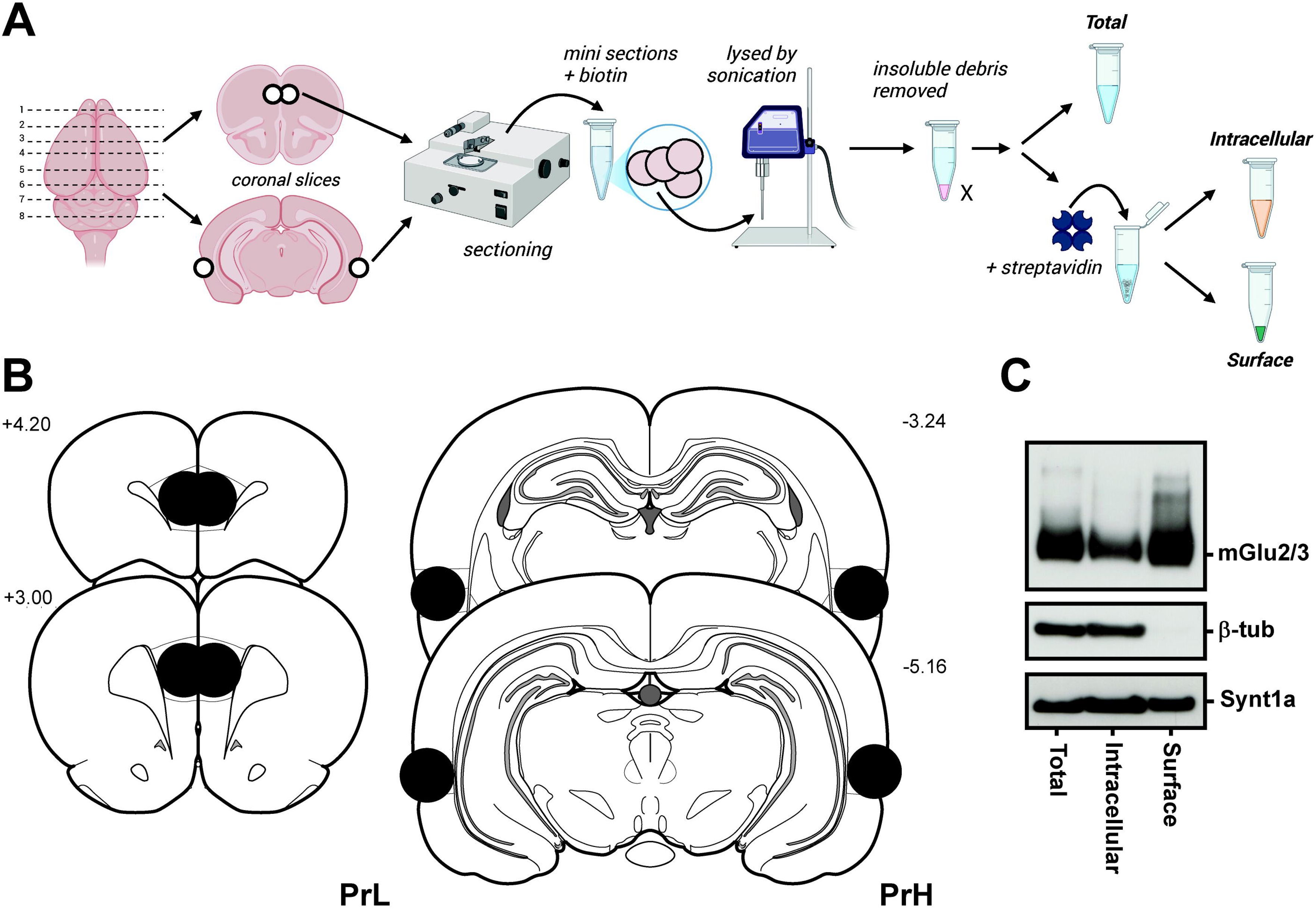
Tissue analysis workflow and representative immunoblotting data. (a) Coronal slices containing the prelimbic cortex (PrL) and perirhinal cortex (PrH) were sectioned into mini-slices, surface-biotinylated, and lysed by sonication. After removal of insoluble debris, total protein was collected, and biotinylated proteins were captured using Neutravidin beads, separating intracellular and surface fractions, subsequently used for immunoblotting analysis; (b) Schematic representation of coronal brain sections indicating the locations of bilateral tissue sampling of the PrL and PrH. The numbers indicate the approximate anterior-posterior distance (in mm) from Bregma; (c) Representative immunoblots showing mGlu2/3 in total, intracellular, and surface fractions. Specific loading controls, β-tubulin (β-tub) and Syntaxin-1a (Synt1a), were used to verify the purity of the intracellular and surface fractions.

### Protein analysis

Because performance on the OIP task depends on coordinated activation of the PrL and PrH, the analysis of protein levels of mGlu2/3 and related proteins focused on these two cortical subregions. Immunoreactivity (in total lysates and in the cell surface fraction) in both subregions was analyzed 24 h after the OIP test. Acute brain slices (containing PrL or PrH) were collected and processed for cell surface biotinylation (as summarized in Figure 4a and in the Methods section). The anatomical location and coordinates depicting bilateral sites of PrL and PrH tissue collection are shown in Figure 4b.

Representative results demonstrating specific surface biotinylation and tissue fractionation are shown in Figure 4c. mGlu2/3 immunoreactivity was detected (as expected) across all fractions, confirming successful protein recovery. β-Tubulin (part of the intracellular cytoskeleton) was present in the total and intracellular fractions but was largely absent from the surface fraction, indicating minimal contamination of the surface fraction by intracellular proteins. Syntaxin-1a (a plasma membrane and presynaptic vesicle regulatory protein) was detected in all fractions and was used to normalize the surface biotinylation of the membrane proteins (namely, mGlu2/3).

In the experimental tissues, protein levels of mGlu2/3, RanBP9, and Gαi (and loading control β-Tubulin) were analyzed in total cellular lysates. In parallel, mGlu2/3 (and membrane loading control Syntaxin-1a) were assessed in the biotinylated cell surface fraction. In the PrL, analysis of the total cell lysate revealed no significant differences between groups for any of the proteins examined, including mGluR2/3 (t(20) = 0.3689, p > 0.05), Gαi (t(21) = 0.9774, p > 0.05), and RanBP9 (t(21) = 0.0079, p > 0.05) (Figure 5a). In contrast, mGlu2/3 levels in the surface-biotinylated fraction obtained from the PrL were significantly higher in the meth group relative to controls (t(21) = 4.413, p < 0.0005) (Figure 5b). Similarly, in the PrH, a history of meth IVSA did not alter total protein levels of either mGlu2/3 (t(18) = 1.005, p > 0.05), Gαi (t(18) = 1.376, p > 0.05), or RanBP9 (t(18) = 0.6686, p > 0.05) compared to controls. Akin to PrL, cell surface levels of mGlu2/3 in the PrH were increased in the meth group, relative to controls (t(18) = 5.791, p < 0.0001) (Figure 5d).

**Figure 5.**
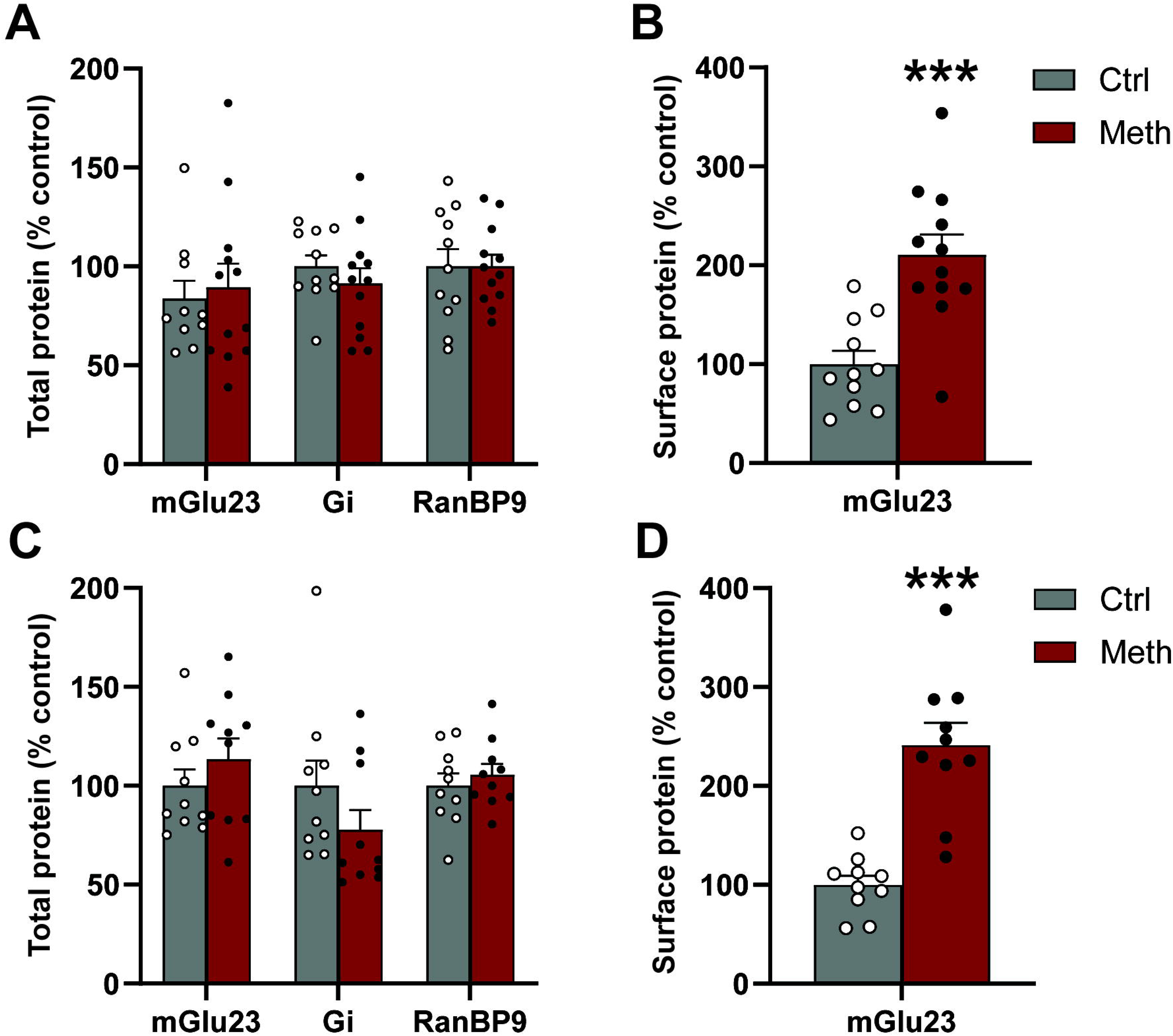
Immunoblotting analysis of total and surface proteins in the PrL and PrH. (a-b) Prelimbic cortex (PrL); (a) Total protein expression of mGlu2/3, Gi, and RanBP9 did not differ between Ctrl and Meth groups (unpaired t-tests: mGlu2/3, t(20) = 0.3689, p > 0.05; Gi, t(21) = 0.9774, p = 0.5014; RanBP9, t(21) = 0.0079, p > 0.05); (b) In contrast, surface expression of mGlu2/3 was significantly increased following meth administration (unpaired t-test, t(21) = 4.413, ***p < 0.0005); (c–d) Perirhinal cortex (PrH). (c) Total protein expression of mGlu2/3, Gi, and RanBP9 was not significantly altered (unpaired t-tests: mGlu2/3, t(18) = 1.005 p > 0.05; Gi, t(18) = 1.376 p > 0.05; RanBP9, t(18) = 0.6686, p > 0.05); (d) However, meth significantly increased surface mGlu2/3 levels (unpaired t-test, t(18) = 5.791, ***p < 0.0001). Groups: Ctr (n = 10-12), Meth (n = 10-12). Data represent the mean ± SEM.

As tissue samples in the meth group originated from rats previously treated with either LY-487379 or vehicle, an exploratory analysis that tested the hypothesis that LY-treatment could have contributed to upregulation of the surface mGlu2/3 signal was conducted. One-way ANOVA revealed a significant effect of treatment on surface mGlu2/3 levels in both the PrL and PrH. In the PrL, surface mGlu2/3 levels differed significantly between groups, F 2, 20 = 12.18, p > 0.001. Tukey’s post hoc comparisons revealed that both meth-LY rats and meth-vehicle rats had significantly higher surface mGlu2/3 levels compared to (vehicle-treated) controls (p < 0.001 and p < 0.05, respectively). However, surface mGlu2/3 levels did not differ significantly between meth-LY and meth-vehicle rats, suggesting that LY-487379 treatment did not further alter the effect of meth. In the PrH, one-way ANOVA also revealed a significant effect of treatment on surface mGlu2/3 levels, F(2, 17) = 28.90, p < 0.0001. Tukey’s post hoc analysis showed that both meth-LY and meth-vehicle rats had significantly higher surface mGlu2/3 levels than controls (p < 0.0001 and p < 0.01, respectively). Importantly, meth-LY rats also showed significantly higher surface mGlu2/3 levels than meth-vehicle rats (p < 0.05), indicating that prior LY-487379 treatment may have further increased surface mGlu2/3 expression in this brain region. Data not graphed.

## 4. Discussion

Here, we describe that extended access to meth in rats produced an escalation of meth intake during IVSA, and impairment in object-in-place recognition memory that persisted up to 30 days into abstinence. Next, we found that systemic administration of mGlu2 receptor PAM (LY-487379) reversed post-meth recognition memory deficit. And finally, immunoblot analysis of mGlu2/3 receptors and associated proteins revealed an increase in mGlu2/3 surface expression, but not total expression, in both PrL and PrH cortices of rats with a history of extended access to meth self-administration.

Repeated meth misuse is associated with an increased risk of developing compulsive, uncontrollable meth seeking and use, leading to MUD. This transition is frequently accompanied by an escalation of meth consumption, and a pattern of prolonged heavy ‘binge’ or ‘run’ use over days or even a week [51,52]. While the escalation of meth seeking and use has been attributed to both the development of tolerance and incentive sensitization, it has been of interest to preclinical research to develop animal models of escalated meth use not only to mimic ‘real-world’ human use patterns, but also to study underlying neural mechanisms and long-term consequences [53,54]. One of the models relies on allowing animal access to intravenous meth during extended (long) self-administration sessions. When compared to limited (1-2 h) access, extended access to meth self-administration (6-12 h/day), (1) approximates the pharmacokinetic pattern of meth abuse observed in humans, (2) predictably escalates meth intake, and (3) produces persistent neurophysiological and behavioral deficits [26,27,55–58]. Here, we found that extended (6h) daily access to meth self-administration produces escalation of intake, reproducing our previous observations [21,26,34,37,57].

Previous work by our lab and others has demonstrated that extended (but not limited) access to meth produces a spectrum of cognitive deficits that coincide with altered synaptic activity and plasticity in the PrL and PrH cortices (see: [59,60]). And further, that selective activation of the PrH-to-PrL circuit rescues meth-induced recognition memory deficits [25]. Specifically, extended access to meth self-administration resulted in early changes in spontaneous neuronal activity, followed by delayed-onset changes in glutamatergic synaptic activity (decrease in the AMPA/NMDA ratio) in the PrL/mPFC [27,58,61], and also impaired long-term depression (LTD) in the PrH [26]. Using the identical self-administration paradigm, extended access to meth produced deficits in object recognition memory [20,21,26], as well as early-onset deficits in attentional processing and cognitive flexibility (reversal learning) [58,62]. One limitation of the above-mentioned studies is that behavioral and neurophysiological changes were detected during early abstinence (within the first 2 weeks of meth discontinuation).

Meanwhile, studies in meth-dependent subjects showed that cognitive deficits (in decision making) and meth craving initially worsen over the first 3 months of abstinence, before recovering [9]. The results presented here show, for the first time, that recognition memory deficits (in the OIP task) persist beyond the first week [20] and remain detectable on day 30 of abstinence in rats with a history of extended access to meth.

The current findings are significant in light of clinical reports suggesting that episodic memory is one of the most consistently and severely impaired cognitive domains in MUD [7,17]. Still, the molecular, cellular, and circuitry maladaptations underlying episodic memory deficits after meth remain incompletely understood.

Dysregulation of multiple neurotransmitter systems—including monoamines, endogenous opioids, and endocannabinoids—has been implicated in meth-associated memory deficits [36,63–65]. However, converging evidence from our laboratory indicates that glutamatergic dysfunction plays a critical role in meth-induced recognition memory deficits. This includes the observation that extended access to meth self-administration reduces mGlu5 receptor expression in the PrH and produces NOR deficits that are rescued by systemic administration of mGlu5 allosteric activator [21]. Another study by our laboratory found that meth-induced NOR deficits can be reversed by direct intra-PrH administration of N-methyl-D-aspartate receptor partial agonist (D-cycloserine) [26]. This study also provided a mechanistic insight into post-meth recognition memory deficits, as it showed that a history of meth disrupts PrH synaptic plasticity (long-term depression, LTD), and that the pro-cognitive effects of D-cycloserine depend on rescuing LTD. With regards to mGlu2/3, our studies found no changes in the total levels of mGlu2/3 in the PFC and PrH after 14 days (cite) or 30 days (current study) of abstinence from extended access meth self-administration. In contrast, two studies in mice found an increase in total cellular levels of mGlu2/3 in the PFC (with no change in the PrH) after a non-contingent low- or high-dose meth administration paradigm at 7 and 25 days of abstinence [35,36]. We hypothesize that several factors may have contributed to the emergence of such conflicting findings. Besides the evident methodological factors, such as the use of distinct model organisms (mouse vs. rat) and distinct meth exposure paradigms (experimenter-administration vs. self-administration), other variables warrant consideration. Given the central role of mGlu2/3 receptors in regulating presynaptic glutamate release, it is reasonable to predict that total mGlu2/3 expression changes gradually over prolonged meth abstinence, paralleling alterations in cortical glutamate content reported in both human subjects and rodent models [66–68]. As such, the initial global decrease in cortical (and subcortical) glutamate levels may be accompanied by decreased mGlu2/3 expression, whereas subsequent glutamate recovery may be complemented by a homeostatic recovery/increase in mGlu2/3 receptor numbers.

Therefore, future studies need to rigorously examine time-dependent changes in mGlu2/3 receptors alongside measurements of glutamate release in the key brain circuits (e.g., cortico-striatal, cortico-perirhinal, and cortico-hippocampal). Surface mGlu2/3 receptors may be subject to additional types of regulation, including neural activity-dependent changes in receptor trafficking [69]. Consistent with this, we previously observed reduced mPFC surface mGlu2/3 expression 14 days after meth, whereas the current study detected increased surface mGlu2/3 in the PrL and PrH after 30 days of abstinence. However, because all animals underwent OIP testing and vehicle/LY administration 24 h before tissue collection, we cannot exclude acute effects of these manipulations on receptor trafficking, particularly in meth-experienced animals (see the Results section for an exploratory analysis of vehicle/LY effects on mGlu2/3 surface expression).

Interestingly, administration of mGlu2 PAM LY-487379 rescued meth-associated deficits in OIP recognition memory in the current study. This finding is in line with prior reports demonstrating the pro-cognitive effects of this PAM on attentional processing and cognitive flexibility in drug-naïve rats, as well as LY’s ability to prevent MK-801-induced disruptions in the NOR and Morris Water Maze memory tasks [60,70,71]. The specific neural mechanism underlying this effect is currently poorly understood. Our earlier study found that meth-associated disruption of LTD in the PrH coincided with NOR recognition memory deficits, and that NMDA receptor activation reversed both LTD and memory deficit [26]. While mGlu2/3-dependent LTD was described in several brain regions, the ability of mGlu2 to regulate LTD in the PrH is less clear. Limited evidence suggests that activation of mGlu2/3 receptors in the PrH is required for the induction of LTD, evoked by mGlu1/5 and NMDA co-activation [72,73]. And while mGlu2/3-independent forms of LTD have also been described across the cortex [74], we hypothesize that LY-487379 activates mGlu2s, restores LTD in the PrH, and rescues meth-associated OIP recognition memory deficits, as observed with D-cycloserine administration [26].

## 5. Conclusions

The data presented here demonstrate that extended access to meth self-administration produces enduring deficits in episodic-like recognition memory that can be rescued by systemic treatment with mGlu2 receptor PAM LY-487379. These findings indicate the utility of mGlu2 allosteric activators for the treatment of meth-associated cognitive deficits. Moreover, as emerging evidence suggests that mGlu2 PAMs can reduce motivation to seek meth and meth reinstatement [75], clinically-relevant mGlu2 PAMs could confer dual benefits in the treatment of MUD. Although orthosteric activation of mGlu2/3 receptors has also been shown to reduce meth-induced hyperactivity and reinstatement of meth seeking [76,77], mGlu2 PAMs are predicted to exhibit superior safety and efficacy in treating SUD [78]. And while these findings are encouraging, significant gaps remain in our understanding of mGlu2/3-mediated mechanisms underlying meth seeking and meth-associated cognitive deficits. Future studies should address additional variables, such as the role of pre-existing cognitive impairment, the contribution of sex differences, and temporal- and circuit-wide adaptations in mGlu2/3 expression and function. In summary, the present findings add to a growing body of evidence implicating mGlu receptors in the neurobehavioral adaptations associated with chronic meth exposure. (for review see: [59]).

## Author Contributions

Conceptualization, M.S.; methodology, M.S., V.G., and L.W.; formal analysis, M.S. and V.G.; investigation, M.S. and L.W.; reagents, M.S. and L.W.; writing—original draft preparation, M.S. and V.G.; writing—review and editing, M.S. and V.G. All authors have read and agreed to the published version of the manuscript.

## Funding

This research was funded in part by grant number R03DA049212 (MS) from the National Institute on Drug Abuse/NIH and by NARSAD Young Investigator Award (MS).

## Institutional Review Board Statement

All animal procedures were conducted at the Medical University of South Carolina (MUSC) between October 2011 and March 2012. The procedures were approved by the MUSC Institutional Animal Care and Use Committee (IACUC) under study protocol no. 2885 (approved in November 2009). All procedures complied with the Guide for the Care and Use of Laboratory Animals (National Research Council (US) Committee for the Update of the Guide for the Care and Use of Laboratory Animals 2011).

## Data Availability Statement

The data presented in this study are deposited in LabArchives and will be made available upon request from the corresponding author due to institutional site license restrictions that prohibit public sharing of web links.

## Acknowledgments

The authors would like to thank Jenna Bilodeau and Rebecca Madell for their technical assistance. The authors would also like to thank Dr. Carmela Reichel for her technical assistance and invaluable advice on experimental design and analysis of the OIP task data. ChatGPT was used to create Graphical Abstract, and Grammarly was used to check grammar. Conflicts of Interest: The authors declare no conflicts of interest. The funders had no role in the design of the study; in the collection, analyses, or interpretation of data; in the writing of the manuscript; or in the decision to publish the results.

## Notes

### Competing Interest Statement

The authors have declared no competing interest.

